# Updating TCGA glioma classification through integration of molecular profiling data following the 2016 and 2021 WHO guidelines

**DOI:** 10.1101/2023.02.19.529134

**Authors:** Mónica Leiria de Mendonça, Roberta Coletti, Céline S. Gonçalves, Eduarda P. Martins, Bruno M. Costa, Susana Vinga, Marta B. Lopes

## Abstract

The understanding of glioma disease has been evolving drastically with dedicated research into the genetic and molecular profiling of glioma tumour tissue. Molecular biomarkers have gained progressive and substantial importance in providing diagnostic information, leading to groundbreaking changes in the tumour classification system, criteria and taxonomy standardised by the 2016 and 2021 editions of the World Health Organization Classification of Tumours of the Central Nervous System’s guidelines (WHO-2016 and WHO-2021, respectively). Some of the insights into glioma disease derived from extensive research on open-source multi-omics databases, such as the Cancer Genome Atlas (TCGA). However, given the substantial changes in glioma classification, retrospective databases may harbour outdated diagnostic annotations, suboptimal for further research. Here we propose two methods for updating the tumour classification of TCGA glioma samples in accordance with WHO-2016 and WHO-2021 guidelines through the integration of curated molecular profiling information. Our methods allowed for the diagnostic update of 98% and 87% of evaluated TCGA glioma cases according to WHO-2016 and -2021, respectively, and highlighted changes in patient-specific diagnosis across both guidelines’ editions. Our reclassification pipelines are provided in software R, facilitating direct reproduction or tailoring upon new releases of WHO guidelines.

## Background

Glioma is the most common form of primary central nervous system (CNS) tumour in adults. These tumours typically exhibit diffuse and infiltrative behaviour, affecting the surrounding tissues and frequently causing disabling or fatal effects (1).

Diffuse glioma pathologic diagnosis has traditionally been based on morphologic (histologic) features. Tumour population cell types were identified (astrocytes, oligodendrocytes or oligoastrocytic/mixed) and assigned a malignancy grade (WHO glioma grade: II to IV) based on the degree of cell proliferation and the presence or absence of necrosis (2). These criteria lead to the recognition of different glioma tumour types, namely *astrocytic tumours* (such as diffuse astrocytoma, anaplastic astrocytoma, and glioblastoma (GBM)), *oligodendroglial tumours* (oligodendroglioma or anaplastic oligodendroglioma) and *oligoastrocytic tumours* (oligoastrocytoma or anaplastic astrocytoma). Tumours could also be described as Lower Grade Gliomas (LGG) according to their lower proliferative activity (WHO grades I, II, and III), excluding the particular case of a diffuse astrocytoma WHO grade IV, denominated as GBM, typically associated with rapid pre- and postoperative disease evolution and fatal outcome (3).

The accelerated knowledge gained about the biology of glioma tumours has led to substantial changes in this disease’s classification over the years, breaking the century-old histogenetic classification of glioma tumours (4, 5) as a result. Particularly, studies on molecular profiling of adult diffuse gliomas have established new insights for the identification of three molecular glioma groups (6–8), first coined by a study by the Cancer Genome Atlas (TCGA) (9): i) “IDH-mutant, 1p/19q codeleted”; ii) “IDH-mutant, 1p/19q non-codeleted”; and iii) “IDH-wildtype”. IDH stands for isocitrate dehydrogenase genes *IDH1* and *IDH2*, and 1p/19q codeletion (1p/19q codel) refers to the concomitant deletion of the 1p and the 19q chromosome arms (and 1p/19q non-codeletion (1p/19q non-codel) otherwise). The terms “IDH-mutant” and “IDH-wildtype” describe, respectively, the presence or absence of mutation in the *IDH1* and/or *IDH2* genes.

Given that the three above-mentioned tumour groups — defined by well-established, simple, and widely available markers — exhibited diverse clinical presentations and different survival patterns (6), a revision of the gold-standard World Health Organization (WHO) classification of gliomas was pushed forward. Thus, with the release of the 2016 WHO Classification of Tumours of the CNS (WHO-2016) (10), glioma classification was refined. For the first time, genetic and molecular profile information became integrated with histologic evaluation, therefore defining or fine-tuning the diagnosis of the disease.

The tendency for reliance on molecular and/or genetic alterations continued to increase in the diagnosis and classification of gliomas. In 2021, the WHO Classification of Tumours of the CNS (WHO-2021) guidelines (11) were published, greatly emphasising the role of genetic markers and molecular profiles of CNS tumours to determine the final diagnosis and to convey prognosis (12). Figure 1 summarises the highlights of the evolution of glioma classification as standardised by the WHO CNS guidelines, highlighting differences and novelties across the most recent versions.

**Figure 1.**
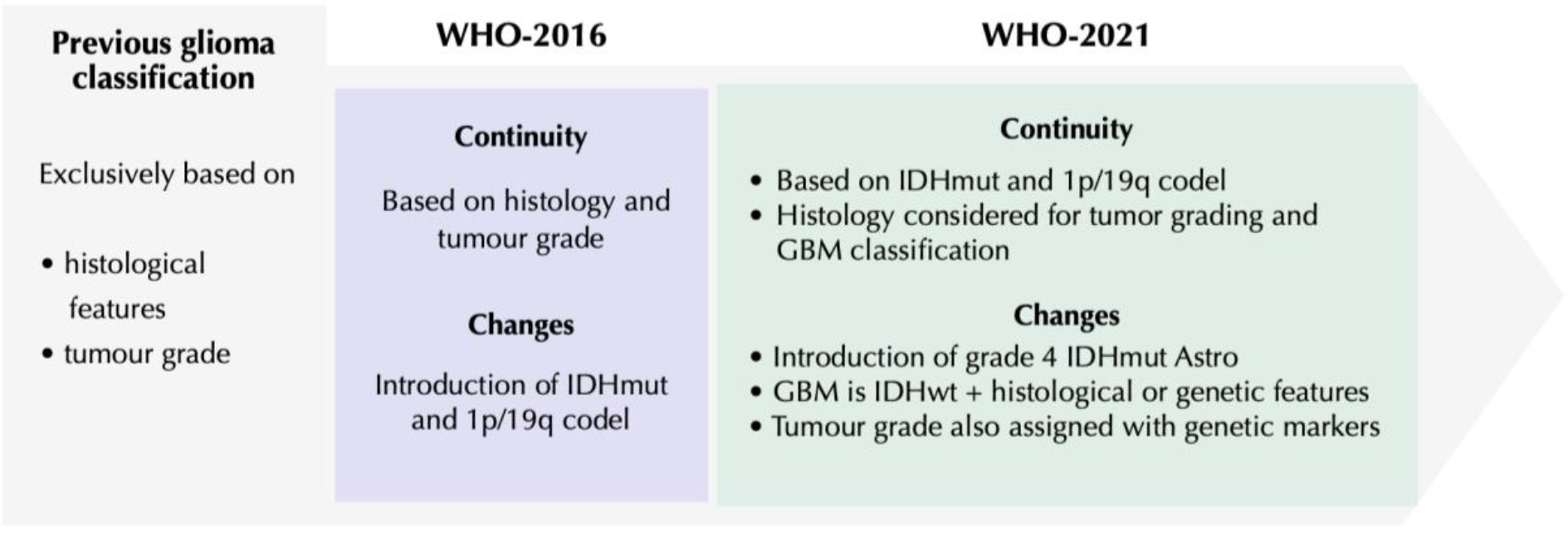
Evolution of WHO CNS diagnostic criteria for glioma tumours before and after WHO-2016 and WHO-2021 releases. The scheme highlights the most prominent differences and similarities between the most recent versions of WHO CNS glioma classification. *Abbreviations: Astro, astrocytoma; codel, codeletion; IDH, isocitrate dehydrogenase; GBM, glioblastoma; mut, mutant; wt, wildtype; WHO, World Health Organization*.

The achieved insights derived from glioma molecular characterisation have profited from the availability of publicly open-source datasets, such as the ones generated by the TCGA program (13). TCGA focused on various cancer types, including gliomas, dedicating specific projects to studying LGG (WHO grades II and III) (14) and GBM (WHO grade IV) (15). However, despite TCGA data being continuously revised and managed by specialised entities (16), its sample data collection ceased in 2013 (17), and some glioma cases were assigned with histological subtypes that are currently known to be strongly discouraged in clinical practice, such as the “mixed glioma” (or oligoastrocytoma) subtype (18). Hence, the diagnostic categories displayed in the TCGA-LGG and - GBM projects’ datasets are not entirely consistent with the taxonomy suggested by WHO-2016 nor WHO-2021. Very recently, Zakharova et al. (19) proposed a procedure for the reclassification of TCGA glioma samples based on WHO-2021 guidelines, yet including molecular features partly differing from the one brought forward with our study. A detailed comparison between our Method-2021 and the referred study is provided in Section B of Supplementary Materials.

Thus, in this work, we propose two methodologies — Method-2016 and Method-2021 — to reevaluate outdated diagnostic annotations of adult glioma samples and, through the integration of curated molecular profiling data and following the WHO-2016 and the WHO-2021 guidelines, assign them with updated tumour classifications. Both methods were applied to the annotated samples belonging to the TCGA-LGG and TCGA-GBM datasets (hereafter referred to as the TCGA-PanGlioma dataset) by integrating correspondent molecular profiling data for each sample. The ultimate goal of both these methods was to divide samples into “simplified tumour classes” of Astrocytoma, Oligodendroglioma and Glioblastoma, so as to facilitate comparison between WHO-2016 and WHO-2021 tumour entities.

An R script (available at https://github.com/RobertaColetti/Update-TCGA-glioma-WHOclassification) was developed to compute the two procedures with the TCGA-PanGlioma dataset and molecular profiling information, giving as an output a final database with glioma tumour types consistent with WHO-2016 and -2021 guidelines, respectively.

### Glioma classifications: 2016 and 2021 WHO CNS guidelines

Here we gather the clinical diagnosis criteria for glioma disease according to the WHO-2016 and -2021. Figure 2 presents a simplification of the process of classification of glioma tumour samples.

**Figure 2.**
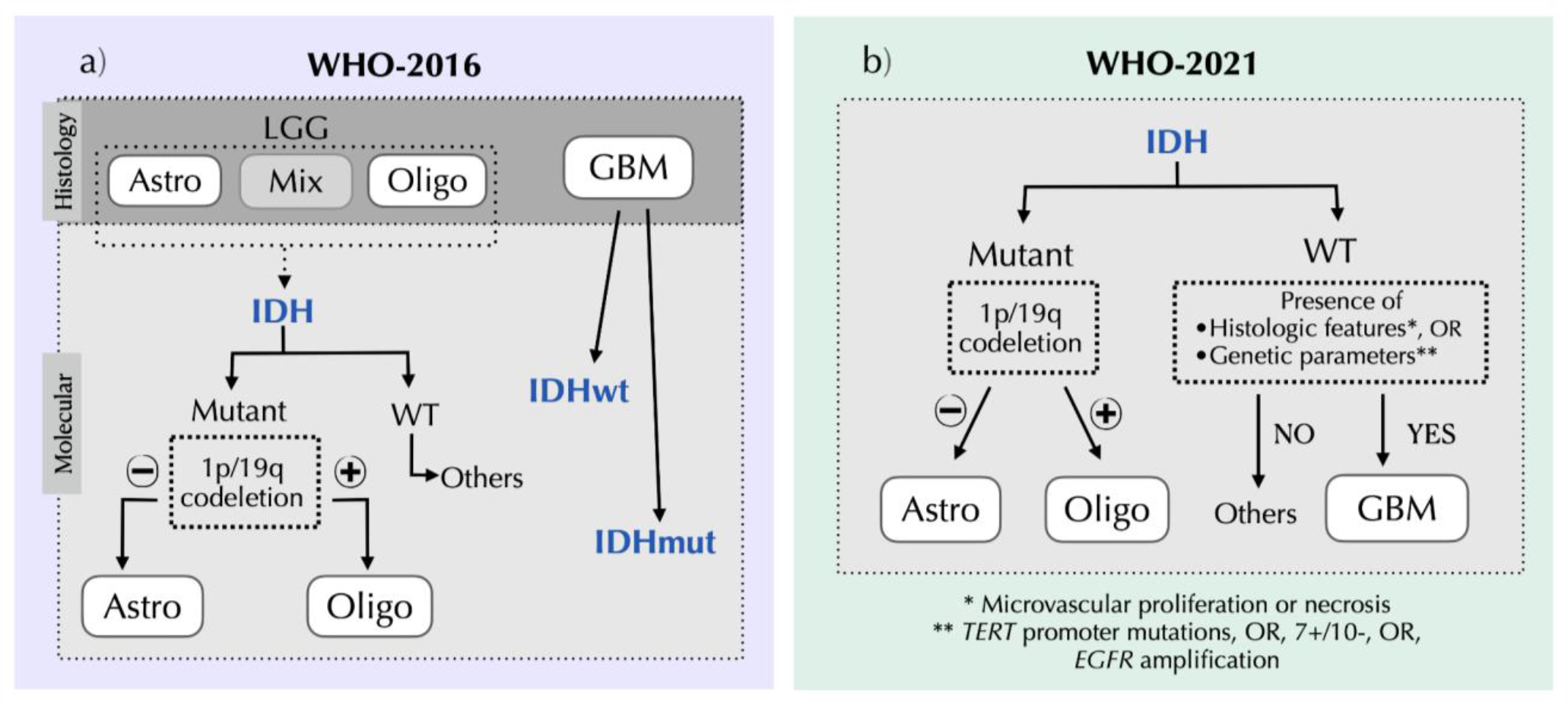
Simplified diagram illustrating the adult glioma classification procedures based on the a) WHO-2016 (Method-2016) and b) WHO-2021 (Method-2021) guidelines. White squares identify the simplified glioma types: astrocytoma (Astro), oligodendroglioma (Oligo) and glioblastoma (GBM). The IDH mutation status (highlighted in blue) can be mutant (mut) or wildtype (wt). There are patients with IDHwt who would require further evaluation to be fully classified; thus we refer to these as “Others’’ in the schemes. **(a)** According to histology, glioma samples were classified as Astro, Oligo, Mixed glioma (Mix) or GBM. Following the WHO-2016 guidelines, samples can be further categorised according to molecular features. In particular, IDHmut LGG samples are classified as Oligo in the presence of 1p/19q codeletion, or classified as Astro in the absence of 1p/19q codeletion. IDHwt LGG samples can be classified as other glioma types (“Others” in the scheme). GBM can be classified as IDHmut or IDHwt. **(b)** According to the WHO-2021 guidelines, the first stage of sample classification is based on their molecular profile. IDHmut can be labelled as Oligo or Astro based on the presence or absence of 1p/19q codeletion, respectively. IDHwt samples that exhibit certain histological/genetic features are classified as GBM. IDHwt samples that do not show these additional features can be considered as other glioma types (“Others” in the scheme). *Abbreviations: Astro, Astrocytoma; [EGFR, epidermal growth factor receptor](https://www.ncbi.nlm.nih.gov/gene/1956); GBM, glioblastoma; IDH, isocitrate dehydrogenase; Mix, Mixed-glioma (i*.*e. oligoastrocytoma); mut, mutant; Oligo, oligodendroglioma; TERT, Telomerase reverse transcriptase; WHO, World Health Organization; wt/WT, wildtype*.

#### 2016 WHO CNS guidelines

The WHO-2016 guidelines presented a major restructuring of the diffuse glioma classification, officially introducing, for the first time, the concept of an “integrated” diagnosis, i.e. combined tissue-based histological and molecular diagnosis (10). This newly introduced diagnostic objectivity aimed at establishing more biologically homogeneous and narrowly defined diagnostic entities than in prior classifications, to improve diagnostic accuracy and patient management, and convey more accurate determinations of prognosis and treatment response (18). Another clear example of the improvements deriving from the changes proposed in WHO-2016 relates to the diagnosis of oligoastrocytoma (or mixed-glioma) — a diagnostic category difficult to define and that suffered from high interobserver discordance and inter-centre variability (20). According to WHO-2016, this category became strongly discouraged in clinical practice, given that nearly all tumours with histological features suggesting both an astrocytic *and* an oligodendroglial component (i.e. “mixed” type) can be classified as either astrocytoma or oligodendroglioma using genetic testing (IDH mutation and 1p/19q codeletion statuses) (18). As a result of the updated categorisation in 2016, both astrocytoma and oligodendroglioma cases became more homogeneously defined (18). Rare cases of IDH-wildtype mixed-gliomas can fall into different unconventional categories (represented in Figure 2a as “Others”).

Thus, regarding glioma tumour classification, WHO grade II diffuse astrocytomas, WHO grade III anaplastic astrocytomas and glioblastomas became divided and further categorised through evaluation of IDH mutation status and/or 1p/19q codeletion. Thus, a tumour was categorised as *IDH-mutant* if exhibiting an IDH mutation or, on the contrary, as *IDH-wildtype*. The designation of *NOS* (Not Otherwise Specified) became reserved for tumours for which full IDH evaluation could not be performed. The final diagnostic nomenclature includes both histologic and molecular profiling evidence (e.g. Diffuse *astrocytoma, IDH-mutant)*, and the grading is predetermined by the diagnosed tumour entity (18).

However, the combined use of histology and molecular genetic features for diagnosis has also created groups of tumours that did not fit into such precise entities. WHO-2016 criteria can possibly lead to uncommon diagnosis, requiring further careful evaluation to avoid misdiagnosis (e.g. diffuse astrocytoma, IDH-wildtype; or anaplastic astrocytoma, IDH-wildtype) (18). The new criteria could also raise discordant tumour-type results. In the latter case, a “two-layered” diagnosis needed to be reported to convey information about findings on both the genotype and histological phenotype (18). For instance, a case of a diffuse glioma with astrocytic phenotype, yet exhibiting IDH mutation *and* 1p/19q codeletion, would necessarily be assigned with two diagnostic annotations: “diffuse astrocytoma, IDH-mutant” (histological phenotype-based diagnosis) and “oligodendroglioma, IDH-mutant and 1p/19q codeleted” (genotype-based diagnosis).

#### 2021 WHO CNS guidelines

The edition of the WHO Classification of Tumours of the CNS published in 2021 (11) builds on the WHO-2016. It establishes some different approaches to both CNS tumour nomenclature and grading, emphasising the role of molecular diagnostics for an “integrated diagnosis”, and greatly simplifying the classification of adult-type diffuse gliomas.

Generally, the major new features in this edition were the official distinction of paediatric-from adult-type tumours and the new approach to grading. Tumours became graded *within* type (rather than across different tumour types), with the WHO CNS grade scale in Arabic numerals from 1–4 (instead of Roman).

While in WHO-2016 diffuse gliomas were highly divided (a consequence of different tumour grades leading to different tumour names/entities), in WHO-2021, adult-type diffuse gliomas are composed of only three types. Namely, i) “*Oligodendroglioma, IDH-mutant and 1p/19q-codeleted”* (grades 2 or 3) in the case of a diffuse glioma exhibiting a combined IDH mutation and 1p/19q codeletion; ii) “*Astrocytoma, IDH-mutant”* (grades 2, 3 and 4) for an IDH-mutant diffuse astrocytic tumour without 1p/19q codeletion; and iii) **“***Glioblastoma, IDH-wildtype****”*** (grade 4). The latter diagnosis is assigned in the setting of an IDH-wildtype diffuse and astrocytic glioma in the presence of certain genetic parameters (*TERT* promoter mutation, or *EGFR* gene amplification, or combined gain of entire chromosome 7 and loss of entire chromosome 10 (+7/−10)), or if exhibiting histologic hallmarks such as microvascular proliferation or necrosis. In the rare occasion that none of the referred genetic or histologic features is present in an IDH-wildtype diffuse and astrocytic glioma, such results do not allow for a WHO diagnosis, and other types of tumour families are considered (e.g. paediatric tumour-type) (represented as “Others” in Figure 2b).

## Datasets and Methodology for Reclassification of Glioma Samples

### TCGA Pan-Glioma Patient Cohort Characteristics

The latest versions of TCGA-LGG and TCGA-GBM cohorts are composed of 515 and 595 patients, respectively, together composing the TCGA-PanGlioma dataset with a total of 1110 glioma cases. The TCGA-LGG project targeted the study of untreated *lower grade gliomas*, consisting of grades II and III, including 194 astrocytomas, 191 oligodendrogliomas, and 130 oligoastrocytomas/mixed-gliomas (9, 14). TCGA-GBM project started in 2008 (15), with samples selected based on an anatomic pathology diagnosis of GBM (equivalent to an astrocytoma WHO grade IV) (8), and it was expanded in 2013 (7), further characterising 595 GBM tumours.

### Molecular Profiling Data of TCGA-PanGlioma Samples

The molecular profiling data is mainly provided by the study of Ceccarelli et al. (6) (*n* = 960 samples). The study performed a comprehensive multi-platform genomic analysis of adult diffuse II, III and IV glioma cases from TCGA, including DNA methylation profile and whole-genome sequencing data analysis. Among other molecular features, it includes information on IDH mutation and 1p/19q co-deletion statuses, as well as *TERT* promoter mutations and the presence of +7/-10. Samples with missing information about *IDH1* and *IDH2* genes statuses (*n* = 140) have been manually searched on Genomic Data Commons (GDC) Data Portal (16), the platform that currently houses and manages TCGA data.

### Data Extraction and Integration

Clinical data from TCGA-LGG and TCGA-GBM projects, including the assigned diagnosis of each tumoural sample (one per patient) was acquired through the *getFirehoseData* function from *RTCGAToolbox* package (21) from R software (version 3.5.1, https://www.r-project.org). We have downloaded the latest Firehose data release available (2016-01-28), as it coincides with the latest clinical data available at the GDC Data Portal (16).

The molecular profiling data, curated by Ceccarelli et al. (6), was downloaded through the *TCGABiolinks* R package (22–24), using the “PanCancerAtlas_subtypes” function and the keywords “lgg” and “gbm”. This data can also be accessed through the supplementary materials provided in the mentioned article. Among the samples included in this study, 140 samples had no information about IDH status. For such cases, we have manually searched for IDH mutation status on GDC Data Portal (16), finding information for 20 samples.

We have merged the two datasets (diagnostic annotation from TCGA and molecular profiling) by the sample’s TCGA-specific barcode, which is composed of a collection of identifiers. We first selected TCGA barcodes referring to primary tumours (sample code “01”), and then reduced the initial TCGA barcode in order to keep only the first two fields (tissue source site and participant, e.g. TCGA-02-0001).

This data integration led to a final TCGA-PanGlioma dataset comprising 1110 samples (515 TCGA-LGG and 595 TCGA-GBM). 12 unmatched samples were excluded, as they were only present in the molecular profiling data and not in the TCGA-LGG-GBM latest data release.

### Methods Description

For reclassification of TCGA-PanGlioma cases according to the WHO-2016 and -2021 guidelines, we developed two methods: Method-2016 and Method-2021, respectively. Both methods require, [to a greater or lesser extent](https://www.ldoceonline.com/dictionary/to-a-greater-or-lesser-extent), processing the diagnosis specified on TCGA-PanGlioma clinical data and the integrated molecular information.

Given the high complexity of the names of glioma tumour entities in WHO-2016 and -2021 guidelines, our main goal is to provide a simplified label in order to divide samples into the three main tumour classes, namely “Astrocytoma”, “Oligodendroglioma” and “Glioblastoma”. Samples that cannot be classified due to lack of information have been labelled as “Unclassified”.

More detailed annotations (in accordance with WHO CNS taxonomy) are available in the Supplementary files “Output1” and “Output2” (https://github.com/RobertaColetti/Update-TCGA-glioma-WHOclassification). A list of the provided R scripts and supporting files can be found in Section A of Supplementary Materials.

#### Method-2016

Method-2016 aims to update the diagnostic annotations of TCGA-PanGlioma cases in accordance with the WHO-2016 (pipeline in Figure 3.a). It focuses on the reclassification of mixed glioma (or oligoastrocytoma) samples using IDH-mutation and 1p/19q codeletion statuses. Glioblastoma samples were further categorised according to IDH-mutation status.

**Figure 3.**
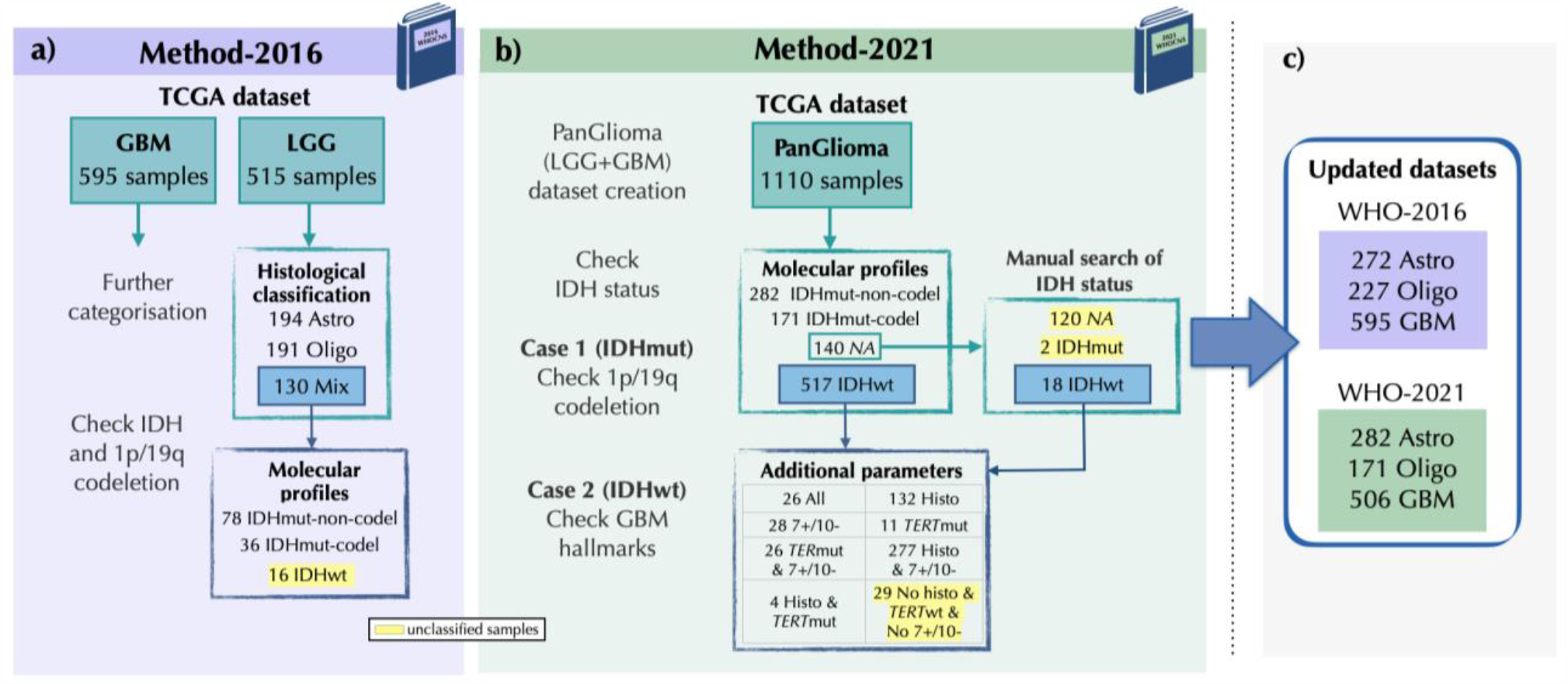
Scheme summarising the classification pipeline of (a) Method-2016 and (b) Method-2021, and the characteristics of the intermediate and final datasets. In the scheme, each step is associated with the number of samples involved in the classification procedure. (c) Size and glioma types of the final updated datasets. *Abbreviations: Astro, Astrocytoma; Chr7+/10-, Chromosome 7 gain / 10 loss; GBM, glioblastoma; Histo, Histologic features; IDH, isocitrate dehydrogenase; LGG, Low Grade Glioma; Mix, Mixed-glioma (i*.*e. oligoastrocytoma); mut, mutant;* NA, *not available; Oligo, oligodendroglioma; TERT, Telomerase reverse transcriptase; WHO, World Health Organization; wt, wildtype*.

Samples with TCGA diagnostic annotation of “anaplastic astrocytoma”, “astrocytoma, NOS”, “Anaplastic oligodendroglioma” or “Oligodendroglioma, NOS” were assigned with a corresponding simplified tumour type label, i.e. either “Astrocytoma” or “Oligodendroglioma”, securing the previous histologic information. This aims to bypass the problematic samples that, according to WHO-2016, would require careful evaluation (see Discussion for more details).

##### i) Reclassification of oligoastrocytoma (mixed glioma) samples

**Step 1:** Select patients with a diagnosis of “mixed glioma” (or, equivalently, “oligoastrocytoma”).

**Step 2:** Consult the IDH status and 1p/19q codeletion status of each patient sample.

**Case 1)** If the patient is “IDH mutant-non-codel”, it is reclassified as “*Astrocytoma*” (*Diffuse astrocytoma, IDH-mutant*).

**Case 2)** If the patient is “IDH mutant-codel”, it is assigned as “*Oligodendroglioma*” (*Oligodendroglioma, IDH-mutant and 1p/19q codeleted*).

**Case 3)** If the patient is “IDH-wildtype”, it is identified as “*Unclassified*”.

##### ii) Further categorisation of Glioblastoma samples

Similarly, we have selected patients with a diagnostic category of glioblastoma. We have evaluated these cases with the integrated information on *IDH* status (wildtype or mutant) from the molecular profiling dataset and assigned them new diagnostic types of either “Glioblastoma, IDH-wildtype” or “Glioblastoma, IDH-mutant” If the sample lacks information on the *IDH* status, it is given the diagnosis of “Glioblastoma, NOS”.

#### Method-2021

With Method-2021 we reclassified each TCGA-PanGlioma case following the WHO-2021 guidelines (pipeline schematised in Figure 3b). We mainly used these patients’ molecular information on “IDH-mutation” and “1p/19q co-deletion” statuses, disregarding the previous clinical annotations from TCGA (derived from histological features). The unique exception regards IDH-wildtype samples, which need additional information to be classified as “Glioblastoma”. In these cases, we considered the status of *TERT* mutation and the presence of +7/-10, as well as the previous TCGA diagnostic annotation. Indeed, the histologic hallmarks used to identify glioblastoma did not change over the years, therefore being classified as “glioblastoma” on TCGA indicates the presence of necrosis or microvascular proliferation. This information, coupled with IDH-wildtype, is sufficient to classify these samples as glioblastoma.

We have considered all samples from the TCGA-PanGlioma dataset. Adopting the WHO-2021 guidelines, we have integrated the clinical and molecular data of 1100 TCGA glioma cases reassigned to the corresponding glioma type according to the following criteria:

**Case 1)** If the sample has IDH mutation:

**Case 1**.**1)** If it has 1p/19q codeletion, it is classified as “Oligodendroglioma” (*Oligodendroglioma, IDH-mutant and 1p/19q codeleted*).

**Case 1**.**2)** If the sample does not exhibit 1p/19q codeletion, it is classified as “Astrocytoma” (*Astrocytoma, IDH-mutant*).

**Case 2)** If the sample is IDH-wildtype:

**Case 2**.**1)** If the TCGA diagnostic annotation was of “glioblastoma” or if a *TERT* promoter mutation is present or exhibits +7/-10, the sample is classified as “Glioblastoma” (*Glioblastoma, IDH-wildtype*).

**Case 2**.**2)** Otherwise, we label the sample as “Unclassified”.

Regarding the samples we integrated manually from TCGA (*n =* 20), 18 belonged to the TCGA-GBM project and were IDH-wildtype, thus diagnosed as “Glioblastoma, IDH-wildtype” according to *Case 2*. The remaining 2 samples were IDH-mutant from the TCGA-GBM project but lacked information on 1p/19q codeletion status to be reclassified according to *Case 1*. These cases were not updated (we could only assess that these 2 samples would transition from GBM to LGG groups).

## Results

The methodology of Method-2016 and -2021 for reevaluation of the original diagnostic annotations of TCGA-PanGlioma samples allowed for the assignment of more updated classes in most cases. Figure 3c reports the dimension of the final updated datasets for each of the methods.

Method-2016 evaluated 130 mixed glioma samples from TCGA-LGG and allowed for the reclassification of 114 cases: 78 astrocytomas and 36 oligodendrogliomas (the remaining samples were unable to be classified due to lack of information to fully follow the guidelines). This method also further characterised 458 glioblastoma samples from TCGA-GBM according to their *IDH* gene mutation status (“Glioblastoma, IDH-mutant” (*n* = 35), “Glioblastoma, IDH-wildtype” (*n* = 423) and “Glioblastoma, NOS (*n* = 137)). This methodology produced an updated version of the TCGA-PanGlioma dataset of 1094 glioma patients’ samples: 272 astrocytomas, 227 oligodendrogliomas and 595 glioblastomas (Figures 3a and 3c).

Method-2021 has evaluated 1110 glioma samples from the TCGA-PanGlioma dataset, generating a final dataset of 959 cases: 282 astrocytomas, 171 oligodendrogliomas and 506 glioblastomas. 151 samples lacked information to be reclassified with this methodology (Figures 3b and 3c).

Table 1 compares the new diagnostic types assigned with Method-2016 and -2021. In the first row, we can assess that 208 (out of 285) samples that were classified as astrocytoma according to Method-2016 maintained this diagnosis with Method-2021, while 5 cases changed to oligodendroglioma, 41 samples changed the classification from astrocytoma to glioblastoma, and 18 were unable to be classified with Method-2021 due to the lack of molecular information. This cross-table highlights the presence of 41 samples changing from oligodendroglioma to astrocytoma, as well as 5 cases that were converted from astrocytoma to oligodendroglioma classes.

**Table 1.** Cross-comparison between the four final diagnostic labels obtained with Method-2016 and with Method-2021: *Astrocytoma, Oligodendroglioma, Glioblastoma* or *Unclassified*. 811 samples (in blue) kept the same classification category on both methods, while 134 cases (in grey) have changed to a different class (unclassified patients not considered).

|  |  | 2021 |  |  |  |
| --- | --- | --- | --- | --- | --- |
|  |  | Astrocytoma | Oligodendroglioma | Glioblastoma | Unclassified |
| 2016 | Astrocytoma | 208 | 5 | 41 | 18 |
|  | Oligodendroglioma | 41 | 164 | 12 | 10 |
|  | Glioblastoma | 33 | 2 | 441 | 119 |
|  | Unclassified | 0 | 0 | 12 | 4 |

Table 1 also emphasises that in the transition from 2016 to 2021 guidelines’ criteria, cases can change from LGG to GBM group, and vice versa. Specifically, 41 2016-astrocytoma and 12 2016-oligodendroglioma samples are respectively classified as 2021-GBM, while 35 2016-GBM would belong to the broader group of LGG (33 astrocytomas and 2 oligodendrogliomas).

## Discussion

CNS tumour classification has been evolving drastically over the years. The stage of understanding of the neuropathology field at a particular time is reflected in every new release of WHO CNS guidelines, providing worldwide practical guidance to pathologists and neuro-oncology specialists and new curated knowledge for the classification of tumour entities.

We have integrated tumour-type annotations of LGG and GBM cases from TCGA with curated molecular profile data, to reevaluate them and reclassify the tumoural samples according to the WHO guidelines of 2016 and 2021 with Method-2016 and -2021. Our reclassification pipelines are easily reproducible through the R scripts (Supplementary Materials, Section A). These scripts can be tailored to other glioma databases, or adapted to cover new diagnostic criteria upon the release of a new version of the WHO guidelines.

Overall, the outcome of the implementation of Method-2016 on TCGA-PanGlioma samples allowed for the reclassification of 114 out of 130 mixed glioma samples, a diagnostic category that became strongly discouraged in clinical practice. Method-2016 was designed to be conservative with respect to the reevaluation of the TCGA diagnosis of astrocytoma, oligodendroglioma and GBM samples (solely based on histology), and exclusively assigning them a univocal simplified glioma subtype label correspondent to the retrospective TCGA diagnosis, in order to secure that same histologic information. Indeed, in WHO-2016, it is warned that this first approach to the usage of integrated diagnosis could possibly lead to the definition of very narrow groups but leaving some glioma cases a bit scattered due to insufficient knowledge about the disease at the time. Thus, our intent with Method-2016 was to avoid assigning fluctuant labels to cases that, according to WHO-2016, would require careful evaluation to avoid misdiagnosis of the glioma lesions. Thus, we bypass the problematic samples that, according to WHO-2016, could lead to: a) two-layered diagnosis or b) tumour entities that were not yet distinguished as solid disease entities at the time (i.e. provisional) (18), or c) uncommon diagnosis (e.g. *diffuse astrocytoma, IDH-wildtype* or *anaplastic astrocytoma, IDH-wildtype*) that would require careful evaluation to avoid misdiagnosis of those glioma lesions. Also, our work highlights such cases when comparing the results from Method-2016 with those of Method-2021, given the improvement of the integrated diagnosis criteria in WHO-2021. In Table 1, we can spotlight the patients that would be assigned with the two-layered diagnosis according to WHO-2016, represented by the cases which changed their WHO-2016 subtype from astrocytoma to oligodendroglioma, according to WHO-2021, and vice versa. This transition underlines the discordance between histological and molecular features that WHO-2016 could give rise to, e.g. samples that are classified as astrocytoma through histological evaluation, yet exhibiting oligodendroglioma molecular features (IDH-mutant and 1p/19q codeleted). Even though the classification provided by Method-2016 does not take into account the complete diagnosis, our procedure avoids introducing uncertainty into the classification for those cases that would need further evaluation. The complexity of this diagnosis has been reduced only with the new WHO-2021 guidelines, which assign a unique label for such cases, exclusively based on genetic and molecular profiles. Specifically, with WHO-2021, the new concept of “integrated diagnosis” has been emphasised, since it combines histological and molecular features to define a univocal final diagnosis, including the tumour grade.

In this work’s Method-2016 and Method-2021, tumour grades were not considered when re-evaluating TCGA-PanGlioma cases. The grading process had undergone significant changes, passing from being tumour entity-specific *to within-tumour-type*, and being influenced by histological features, molecular biomarkers and clinical context (25, 26). The data constraints of this retrospective study impair the tumour grade re-evaluation for the complete TCGA-PanGlioma dataset, which would require sample-specific revision of the histopathological features. Moreover, nowadays the necessity of assigning a grade to every case is still under debate as it may be a source of confusion in clinical care (12). Indeed, currently, CNS tumour grades may no longer reflect the traditionally expected clinical-biological behaviour, given the great impact that modern molecular-based treatments could have on patients’ prognosis (27).

Finally, although the integration of molecular profile data with TCGA diagnostic annotation allowed us to establish WHO CNS diagnostic classes for the great majority of glioma cases, there were samples unable to be classified. While with Method-2016 only 1% of all the cases could not be updated, with Method-2021 the number of unclassified samples substantially increased to 13% (*n* = 151). This has occurred due to: i) overall unavailable IDH status information (*n* = 120), ii) lack of 1p/19q codeletion status data for IDH-mutant cases manually searched on GDC Data Portal (16) (*n* = 2), and iii) IDH-wildtype cases requiring additional information to acquire the diagnosis of “Glioblastoma, IDH-wildtype” (*n* = 29). The latter happens due to the fact that we had no access to one of the three genetic/molecular markers (*EGFR* amplification) that can define this diagnosis.

Despite the retrospective nature of the data from TCGA, the integration of molecular profiling data enabled the update of 1094 (98%) and 959 (87%) glioma types following the latest WHO guidelines of 2016 and 2021, respectively. By spotting cases with outdated diagnostic labels (e.g. “mixed glioma”) and by highlighting the possibility of substantial changes in glioma type and/or their broader categories (LGG/GBM) based on molecular data, our study fosters further data analysis of TCGA-PanGlioma cases. These variations can have a considerable impact on future bioinformatic and statistical analyses that aim at studying and characterising different glioma cases. It is hoped that this pipeline of reclassification of tumour types provides the scientific community with new pathways of analysis of glioma disease from genomic databases, and ultimately benefit patients affected by this disease.

## Author contributions

**Mónica L. Mendonça**: Writing - Original Draft, Conceptualisation, Methodology, Data curation, Visualisation.

**Roberta Coletti:** Conceptualisation, Methodology, Software, Data curation, Visualisation, Writing - Original Draft.

**Céline S. Gonçalves:** Conceptualisation, Validation, Writing - Review & Editing.

**Eduarda P. Martins:** Validation, Writing - Review & Editing.

**Bruno M. Costa:** Conceptualisation, Validation, Writing - Review & Editing, Funding acquisition.

**Susana Vinga:** Supervision, Writing - Review & Editing.

**Marta B. Lopes:** Conceptualisation, Supervision, Writing - Review & Editing, Funding acquisition.

## Acknowledgments

This work was supported by national funds through Fundação para a Ciência e a Tecnologia (FCT) with references CEECINST/00102/2018, 2021.02600.CEECIND, PD/BDE/143154/2019, COVID/BDE/153298/2023, UIDB/00297/2020 and UIDP/00297/2020 (NOVA Math, Center for Mathematics and Applications), UIDB/04516/2020 (NOVA LINCS), UIDB/50026/2020 and UIDP/50026/2020 (ICVS), UIDB/50021/2020 (INESC-ID), UIDB/50022/2020 (IDMEC), and the research project “MONET – Multi-omic networks in gliomas” (PTDC/CCI-BIO/4180/2020). This work was also supported by the project NORTE-01-0145-FEDER-000055, supported by Norte Portugal Regional Operational Programme (NORTE 2020), under the PORTUGAL 2020 Partnership Agreement, through the European Regional Development Fund (ERDF). This project has received funding from the European Union’s Horizon 2020 research and innovation program under Grant Agreement No. 951970 (OLISSIPO project).

Datasets: The results shown here are in whole or part based upon data generated by the TCGA Research Network: https://www.cancer.gov/tcga.

