## Supplementary Materials for "Updating TCGA glioma classification through integration of molecular profiling data following the 2016 and 2021 WHO guidelines"

### A. R programming scripts and supporting files

#### R scripts and files

Codes were written in R (version 3.5.1, https://www.r-project.org) programming language, and supporting files are available at https://github.com/RobertaColetti/Update-TCGA-glioma-WHOclassification. They are organised in two folders named "Scripts-for-reproducibility" and "Final-outputs". The following Table A.1 proves a list of scripts and files included in these folders.

| Scripts | Files | Descriptions |
|---|---|---|
| 2016-classification.R | INPUT<br>• TCGA-LGG.RData<br>• TCGA-GBM.RData<br><br>OUTPUT<br>• **Integrated_data_2016.RData** | TCGA-LGG and -GBM clinical information<br><br>Matrices (one per each glioma type + unclassified) containing clinical + integrated molecular information needed for reclassification |
| 2021-classification.R* | INPUT<br>• TCGA-LGG.RData<br>• TCGA-GBM.RData<br><br>• IDHstatus-TCGA-case_study.csv<br><br>OUTPUT<br>• Case_set.csv<br>• **Integrated_data_2016.RData** | TCGA-LGG and -GBM clinical information<br><br>Information about IDH status retrieved manually<br><br>List of cases with missing IDH status information<br><br>Matrices (one per each glioma type + unclassified) containing clinical + integrated molecular information needed for reclassification |
| Creation_output1.R | INPUT<br>• **Integrated_data_2016.RData**<br>OUTPUT<br>• **Matrix_WHO2016.csv** | Sample classification provided by Method-2016 |
| Creation_output2.R | INPUT<br>• **Integrated_data_2021.RData**<br>OUTPUT<br>• **Matrix_WHO2021.csv** | Sample classification provided by Method-2021 |
| Creation_output3.R | INPUT<br>• **Integrated_data_2016.RData**<br>• **Integrated_data_2021.RData**<br><br>OUTPUT<br>• **SIMPLIFIED_CLASSIFICATION_TCGA_2016_2021.csv** | Comparison between TCGA, 2016 and 2021 classifications (simplified labels) |

**Table A.1.** List of scripts necessary to reproduce Method-2016 and -2021 available at https://github.com/RobertaColetti/Update-TCGA-glioma-WHOclassification. Files in bold are further described in the item below.

* In "2021-classification.R" script, a commented line (No. 84) creates the csv file with the samples we manually searched on the portal ("case_set.csv" file). The file "IDHstatus-TCGA-case_study.csv" needs to be uploaded to include the information we

integrated from the GDC Data Portal. It contains the list of *n* = 20 samples providing the IDH status information, the outcome of our information retrieval (WT or *IDH1* and/or *IDH2*-mutant), and the corresponding mutation, if present.

#### Additional information about files

- **"Integrated_data_2016.RData" and "Integrated_data_2021.RData":** R files comprising four matrices each (one per glioma type – considering the simplified labels: Astrocytoma, Oligodendroglioma and Glioblastoma –, plus a list of the unclassified samples). These matrices contain all the clinical information provided by the TCGA-LGG and -GBM projects, integrated with the curated molecular features we need for reclassification. Particularly, we included IDH mutation status and the presence/absence of 1p/19q codeletion, and, only in the case of Method-2021, information about *TERT* promoter status and the combination of chromosome 7 gain and chromosome 10 loss, to be able to classify glioblastoma in samples exhibiting IDH-wildtype.

- **"Matrix_WHO2016.csv"** (Output 1) resumes the sample classification provided by Method-2016. This 1110 x 4 matrix reports, for each sample, the corresponding histological type currently available on TCGA ("TCGA_subtype" column), the exact type nomenclature assigned by our method, in accordance with the 2016 WHO CNS taxonomy ("new_nomenclature_of_histological_subtype" column), as well as the simplified name we attributed ("simplified_label" column).
  The WHO-2016 nomenclature terms considered were:
  - *diffuse _astrocytoma_IDHmut* (if astrocytoma or mixed glioma with IDH mutation)
  - *oligodendroglioma_IDHmut_codel* (if oligodendroglioma or mixed-glioma with IDH mutation and 1p/19q codeletion)
  - *diffuse_astrocytoma_IDHwt* (if astrocytoma with IDHwt)
  - *oligodendroglioma_NOS* (if oligodendroglioma with either IDHwt or IDHmut but 1p/19q non-codel, or missing information on IDH mutation and 1p/19q codeletion status)
  - *diffuse_astrocytoma_NOS* (if astrocytoma with missing information on IDH status)
  - *glioblastoma_IDHwt* (if glioblastoma with IDHwt)
  - *glioblastoma_IDHmut* (if glioblastoma with IDH mutation)
  - *glioblastoma_NOS* (if glioblastoma with missing information on IDH status)
  - *unclassified* (if mixed glioma with IDHwt)

- **"Matrix_WHO2021.csv"** (Output 2) resumes the sample classification provided by Method-2021. This 1110 x 5 matrix reports, for each sample, the corresponding histological type currently available on TCGA ("TCGA_subtype" column), the exact type nomenclature assigned by our method, in accordance with the 2021 WHO CNS taxonomy ("new_nomenclature_of_histological_subtype" column), as well as the simplified name we attributed ("simplified_label" column). This matrix also contains a "GBM_genetic_parameter" column, which specifies, for each GBM sample, the genetic parameters which allow us to identify the GBM subtype (either *TERT* mutation or Chromosome 7 gain / 10 loss). The WHO-2021 nomenclature terms considered were: "astrocytoma_IDHmut" (if IDHmut without 1p/19q codel), "olgodendroglioma_IDHmut_codel" (if IDHmut with 1p/19q codel), "glioblastoma_IDHwt" (if IDHwt + histological or genetic hallmarks) and "unclassified" (in case of missing information).

- **"SIMPLIFIED_CLASSIFICATION_TCGA_2016_2021.csv"** (Output 3), which compares, for each sample, the current TCGA histological type classes ("TCGA.classification" column), with the simplified label we provide by applying Method-2016 ("classification.2016" column) and Method-2021 ("classification.2021" column).

The last three files are in the "Final-Outputs" folder at https://github.com/RobertaColetti/Update-TCGA-glioma-WHOclassification.

#### B. Comparison between our study's *Method-2021* and the study of Zakharova et al.

A recent study by Zakharova et al. (1) has proposed a procedure for diagnostic reevaluation of TCGA-LGG and -GBM glioma samples based on the WHO-2021 guidelines (2). As in our Method-2021, the authors have integrated data annotation from TCGA-LGG and TCGA-GBM projects and molecular profiling data from the study of Ceccarelli et al. (3), yet including molecular features partly differing from our method.

Our procedure led to the update of 941 out of 1110 glioma cases with a reproducible and automated pipeline, while the procedure of Zakharova et al. led to the update of 828 out of 1122 samples. Comparing the final tumour classes obtained (Table B.1), we conclude that the classification outcomes are in agreement, except for cases defined as "Unclassified". Particularly, our Method-2021 has classified 143 additional samples, representing extra 11% astrocytomas, 12% oligodendrogliomas and 17% GBM cases; while the pipeline of Zakharova et al. has further classified 10 glioblastomas (2% of GBM cases) and 2 astrocytomas (<1% of astrocytoma cases), about which we did not find IDH mutation information.

| | | Our classification (Method-2021) ($n$ = 1110) | | | |
|---|---|---|---|---|---|
| | | **Astrocytoma** | **Oligodendroglioma** | **Glioblastoma** | **Unclassified** |
| Zakharova et al. (1) ($n$ = 1122*) | **Astrocytoma** | 250 | 0 | 0 | 2 |
| | **Oligodendroglioma** | 0 | 150 | 0 | 0 |
| | **Glioblastoma** | 0 | 0 | 416 | 10 |
| | **Unclassified** | 32 | 21 | 90 | 139 |

Table B.1 - Cross-comparison between tumour type classes obtained with Method-2021 and the one proposed by Zakharova et al. (1), including unclassified samples. Yellow cells highlight the samples assigned to the same class by both approaches, in grey cells the discordant tumour classes, and in blue the cases unable to be classified by both studies. Our procedure classified additional 143 cases (32 astrocytomas + 21 oligodendrogliomas + 90 glioblastomas), while Zakharova et al. classified 12 extra cases (2 astrocytoma + 10 glioblastomas). Note: *Only the $n$ = 1110 common samples between the studies were considered (12 extra samples in the initial cohort of Zakharova et al. were excluded by the authors due to unknown histology).

Next, we list and discuss the discrepancies between our Method-2021 and the pipeline proposed by Zakharova et al., in light of current biological knowledge and the 2021 WHO CNS guidelines.

##### I. Tumour grading

The reclassification procedure proposed by Zakharova et al. (1) provides both the tumour type and grade. According to WHO-2021, the grade is one of the diagnostic layers characterising tumour malignancy, and newly recognised tumour types have accepted grades, e.g. a "Glioblastoma, IDH-wildtype" is, by definition, WHO grade 4. Conversely, Astrocytoma and Oligodendroglioma tumours can be associated with different grades, thus grading these two tumour types is done according to specific histologic and/or molecular criteria.

Traditionally, CNS tumour grading has been solely based on histological evaluation. However, the recent WHO-2021 guidelines introduced the usage of biomarkers to complement the process of grade assignment.

Particularly for adult-type diffuse gliomas, WHO-2021 includes evaluation of *CDKN2A*/*B* homozygous deletion (HD) in IDH-mutant astrocytomas. When this biomarker is positive in an astrocytic IDH-mutant tumour, the highest grade is assigned, being diagnosed as "Astrocytoma, IDH-mutant, WHO grade 4".
On the other hand, grading criteria for oligodendroglial tumours remained mainly based on histological features, which should be considered to unequivocally assign WHO grades 2 or 3. Despite this, the WHO-2021 guidelines also mention the evaluation of *CDKN2A* homozygous deletion as a molecular biomarker to assign

oligodendroglioma to WHO grade 3 in particular cases, such as "in tumour samples with borderline histological features" (4).

According to the reclassification scheme of Zakharova et al. (Figure 2 in (1)), the authors have applied the evaluation of *CDKN2A*/*B* HD when assigning grades to either IDH-mutant astrocytomas or oligodendrogliomas, particularly:

a) astrocytomas *with CDKN2A*/*B* HD assigned to the maximum grade 4;
b) oligodendrogliomas *with CDKN2A* HD assigned to the maximum grade 3;
c) astrocytomas *without CDKN2A/B* HD constricted to temporary grades 2 or 3;
d) oligodendrogliomas *without CDKN2A* HD constricted to temporary grades 2 or 3.

While the criterion a) is defined according to the novelty introduced with the latest guideline's releases, the criterion b) could be debatable. Several studies investigating the role of *CDKN*2A in oligodendroglioma prognosis led to inconsistent conclusions (5–9). Moreover, WHO-2021 guidelines mention the usefulness of testing for *CDKN2A* HD in particular scenarios, but not routinely required (4). As a consequence, upgrading an oligodendroglioma sample to grade 3 based on this marker should be performed only after a careful evaluation of the patient-specific clinical profile.

Concerning criteria c) and d), it is not clarified in the manuscript of Zakharova et al. how the final grading (grade 2 or 3) is assigned. According to our interpretation of supplementary materials of the study of Zakharova et al. (Table S1 in (1)), if homozygous deletion is unknown or negative, the samples maintain the retrospective grading annotation (e.g. a case with previous grade II reclassified as astrocytoma, gets labelled as "Astrocytoma, IDH-mutant, grade 2").

The methodology applied by Zakharova et al. led to the update of astrocytoma and oligodendroglioma grades for 45 cases ($n$ = 43 "Astrocytoma, IDH-mutant, grade 4", and $n$ = 2 "Oligodendroglioma, IDH-mutant and 1p/19q-codeleted, grade 3"), while 357 cases maintained the retrospective grades (coinciding with the one provided by Ceccarelli et al. (3)).

Our proposed methodology did not aim to re-evaluate tumour grades (as mentioned in the Discussion section), instead following a more conservative approach and updating the samples with the three main glioma types. One of our motivations lied in the fact that we would be able to do it for a subset of cases (specifically 33 "Astrocytoma, IDH-mutant" would be up-staged to grade 4, when the sample was previously included in the TCGA-GBM project, meaning it necessarily showcased microvascular proliferation or necrosis).

#### II. *ATRX* status

*The ATRX* gene is known to be characteristically altered in IDH-mutant astrocytomas (2). Also, the WHO-2021 guidelines designate the presence of *ATRX* mutation as a sufficient condition to assign IDH-mutant cases to astrocytoma type (4, 10).

In the re-classification method proposed by Zakharova et al., *ATRX* gene status is used to assign an updated diagnosis of "Oligodendroglioma, IDH-mutant, 1p/19q-codeleted". Particularly, based on Zakharova et al. reclassification scheme (Figure 2 in (1)), an IDH-mutant case previously labelled by TCGA as "Glioblastoma", would be classified as "Oligodendroglioma, IDH-mutant, 1p/19q-codeleted" if it exhibited 1p/19q codeletion *or* if it was *ATRX*-wildtype. Even though mutual exclusion between *ATRX* mutation and 1p/19q codeletion has been observed (11), the opposite is not verifiable, i.e. *ATRX*-wildtype does not indicate the presence of 1p/19q codeletion (12). For instance, in Ceccarelli et al. (3) (which both our study and the study of Zakharova et al. used to retrieve the majority of molecular profiling data), 18% of IDH-mutant samples ($n$ = 77 out of 437) do not exhibit such mutual exclusivity, presenting *ATRX*-wildtype without 1p/19q codeletion. Moreover, WHO-2021 guidelines do not establish *ATRX* gene alteration as a definitive diagnostic marker for oligodendroglioma, and the evaluation of 1p/19q codeletion is necessary for its diagnosis (2).

However, a thorough analysis of Table S1 of Zakharova et al. (supplementary material) revealed that the authors had assigned the updated diagnosis of 'Oligodendroglioma, IDH-mutant, and 1p/19q-codeleted' if the sample exhibited both *ATRX*-wildtype and 1p/19q codeleted. Instead, IDH-mutant *and* 1p/19q-codeleted cases with *ATRX* mutation were excluded by the authors, stating that «as "oligodendrogliomas, IDH-mutant, and 1p/19q-codeleted" lack *ATRX* mutation these cases are left unclassified».

In practice, although *ATRX* mutation is sufficient for an IDH-mutant astrocytoma diagnosis, «retained nuclear *ATRX* positivity in an IDH-mutant glioma should prompt analysis for 1p/19q codeletion in order to distinguish IDH-mutant astrocytoma from IDH-mutant and 1p/19q- codeleted oligodendroglioma. *ATRX* immunohistochemistry is not necessary if IDH mutation and 1p/19q codeletion status are captured within one more extensive molecular marker panel assay» (13).
The choice to evaluate *ATRX* status in the study of Zakharova et al. led to the exclusion of *n* = 3 samples (2 TCGA-oligodendrogliomas, 1 TCGA-oligoastrocytoma), even though they present both IDH mutation and 1p/19q codeletion. These samples, according to our Method-2021, were reclassified as Oligodendroglioma.

#### III. H3 status

Overall, the criteria leading to the biggest difference between the outcome from our classification method (Method-2021) and the pipeline of Zakharova et al. (1) is related to the histone 3 alterations (H3 status) for reclassification of a sample into "Glioblastoma, IDH-wildtype".

Our Method-2021 pipeline (Figure 2b, main manuscript) has conservatively followed the diagnostic criteria indicated by the WHO-2021 guidelines, according to which a «Glioblastoma, IDH-wildtype should be diagnosed in the setting of an IDH-wildtype diffuse and astrocytic glioma in adults if there is microvascular proliferation or necrosis or *TERT* promoter mutation or *EGFR* gene amplification or 7+/10- chromosome copy number changes» (2), leading to the reclassification of *n* = 506 glioblastoma samples.

According to Zakharova et al., "Glioblastoma, IDH-wildtype" diagnosis can be assigned to the cases exhibiting IDH-wildtype and H3-wildtype and one of the above-mentioned histological and genetic hallmarks, thus reclassifying *n* = 416 cases.

As reported in the WHO-2021 guidelines, *Glioblastoma, IDH-wildtype* is defined as «a diffuse, astrocytic glioma that is IDH-wildtype and H3-wildtype and has one or more of the following histological or genetic features: microvascular proliferation, necrosis, *TERT* promoter mutation, *EGFR gene amplification, +7/−10 chromosome copy-number changes»* (4). H3-wildtype is a molecular characteristic of most glioblastoma cases, but the diagnosis of adult-type glioblastoma and presence of H3 mutation are not necessarily mutually exclusive (https://doi.org/10.1111/bpa.13062). In fact, while the H3 status plays a substantial role in the paediatric-type glioma classification (2, 14), it is not explicitly mentioned in the primary adult-type glioma diagnostic criteria (2). In clinical practice, testing for H3 status is not necessary to attribute a diagnosis of adult-type glioblastoma, which is diagnosed in presence of IDH-wildtype diffusely infiltrating astrocytic gliomas, showing one of the histological or genetical features mentioned above (2). Conversely, «IDH-wildtype gliomas that do not fulfil these diagnostic criteria should prompt investigation for other possible entities, such as H3-mutant or MAPK driven gliomas» (15).

The evaluation of H3 status in the methodology of Zakharova et al. has led to the exclusion of *n* = 82 IDH-wildtype with *unknown H3*, and *n* = 3 IDH-wt and H3-mutant. Of those, we were able to reclassify *n* = 83 as "Glioblastoma, IDH-wildtype".

**Final considerations**

The role of molecular markers for glioma diagnosis and prognosis has been and continues to be extensively studied (16), with the ultimate goal of achieving better criteria for patient stratification. The process of integration of such findings into clinical practice is extensive and extremely rigorous. The guidelines outline a variety of molecular characteristics that can be essential to establish a diagnosis. However, referring to multiple criteria in a retrospective data analysis could lead to conflicting outcomes, affecting the process of diagnostic label re-assignment. Our approach was to define reclassification criteria for the three glioma types that allowed us to establish a solid and straightforward diagnosis according to WHO-2021.

Moreover, it is worth noting that the reclassification of retrospective open-source data that can be retrieved with different methodologies (e.g. manual download through GDC Data portal (16), automatic data loading through R software packages) may contribute to possible differences in the versions of the collected data and, thus, in the final reclassification outcomes. Thus, further efforts are needed to standardise the reclassification of TCGA samples through the integration of external sources according to the WHO guidelines.
